# RAG-ESM: Improving pretrained protein language models via sequence retrieval

**DOI:** 10.1101/2025.04.02.646805

**Authors:** Damiano Sgarbossa, Anne-Florence Bitbol

## Abstract

Protein language models are significantly advancing the modeling of sequence-function relationships. However, most of them are not directly informed of homology and evolutionary relationships between protein sequences. Here, we propose a method to make them homology-aware. We introduce RAG-ESM, a retrieval-augmented framework that allows to condition pretrained ESM2 protein language models on homologous sequences, using a minimal number of additional cross-attention parameters and minimal computational cost. We show that RAG-ESM models outperform larger ESM2 models for masked amino acid prediction. We find that sequence alignment capabilities spontaneously emerge in specific cross-attention heads of RAG-ESM. By using a discrete diffusion objective for training, and by conditioning on homologs during inference, RAG-ESM reaches state-of-the-art performance for conditional protein sequence generation and motif scaffolding, among sequence-based models. Our method thus possesses strong potential for scalable, efficient and controlled protein engineering.

## 1 Introduction

The three-dimensional structures and biological functions of proteins are encoded in their amino-acid sequences. Homologous proteins share a common ancestry, and have similar structure and function. Natural selection for function gives rise to statistical signatures in their sequences. Homology and evolutionary information are thus extremely useful for modeling the sequence-function relationship, and for protein engineering and mutational effect prediction. Experimental approaches like directed evolution and mutational scanning are usually restricted to the local neighborhood of an existing protein sequence. However, the expansion of large-scale databases, such as UniProt (The UniProt Consortium, 2021), has facilitated computational modeling leveraging evolutionary diversity (Weigt et al., 2009; Morcos et al., 2011; Marks et al., 2011; Russ et al., 2020b; Hawkins-Hooker et al., 2021).

Language models trained on large ensembles of protein sequences produce representations of proteins that correlate with their function (Elnaggar et al., 2021; Vig et al., 2021; Rives et al., 2021; Madani et al., 2020; 2023), and enable sequence generation (Ferruz et al., 2022; Madani et al., 2023) and mutational effect prediction (Meier et al., 2021; Kantroo et al., 2024). These models can be recurrent neural networks (Bepler & Berger, 2019), transformers (Rives et al., 2021) or state space models (Sgarbossa et al., 2024), and are trained with objectives such as masked language modeling, autoregressive generation, or discrete diffusion (Alamdari et al., 2023; Wang et al., 2025). While most of these protein language models (pLMs) were trained on unstructured ensembles of single sequences, some were trained on multiple sequence alignments (MSAs) of homologous proteins, and can thus directly exploit evolutionary diversity and functional constraints. MSA-based pLMs include MSA Transformer (Rao et al., 2021) and AlphaFold2’s EvoFormer (Jumper et al., 2021). Despite being trained only on sequences, MSA Transformer was shown to encode structural contacts between amino acids in its row attention heads (Rao et al., 2021). The training of MSA-based pLMs to fill in masked amino acids using the surrounding context of an MSA allows them to efficiently capture coevolution between amino acids due to structural constraints, with far fewer parameters than models based on single sequences (Rao et al., 2021; Lin et al., 2023). These models are an important ingredient of AlphaFold’s major advance in structure prediction Jumper et al. (2021); Elofsson (2023). However, MSA-based models are memory-intensive (Rao et al., 2021), and may inherit the imperfections of MSAs (Thompson et al., 2011). Recently, autoregressive models have been trained on concatenations of non-aligned homologs (Truong Jr & Bepler, 2024; Sgarbossa et al., 2024). However, starting from long sequences of concatenated homologs poses memory issues for transformer-based models.

We posit that augmenting pretrained single-sequence pLMs by using homology could improve their performance, thereby combining the advantages of single-sequence and MSA-based models. To investigate this, we use Retrieval-Augmented Generation (RAG) to improve the pLM ESM2 (Lin et al., 2023), by using homologous sequences as the retrieved external sources. RAG allows to improve the accuracy of generative large language models (LLMs) by integrating relevant data from external knowledge sources (e.g. domain-specific information) into the generation process (Lewis et al., 2020; Guu et al., 2020). Our model RAG-ESM outperforms substantially larger ESM2 models at masked amino acid prediction. Furthermore, we find that sequence alignment capabilities emerge in cross-attention heads. By using a discrete diffusion objective during training and conditioning on homologs during inference, we perform conditional protein sequence generation and motif scaffolding using RAG-ESM, and obtain state-of-the-art performance. We further show that retrieval yields significant improvements on the structural and evolutionary fidelity of generated sequences. RAG-ESM builds on pretrained pLMs, and improves their performance and efficiency. It thus possesses strong potential for scalable and efficient protein engineering.

## 2 Methods

### 2.1 Summary of our contributions

In this work, we demonstrate that retrieval techniques can improve pretrained single-sequence pLMs, such as ESM2 (Lin et al., 2023). Inspired by Zheng et al. (2023), where pretrained pLMs were conditioned on structural information through a structure encoder and cross attention on the final layer, we similarly condition pretrained pLMs, but using sequence information (see Fig. 1 (left) and Section 2.2).

**Figure 1:**
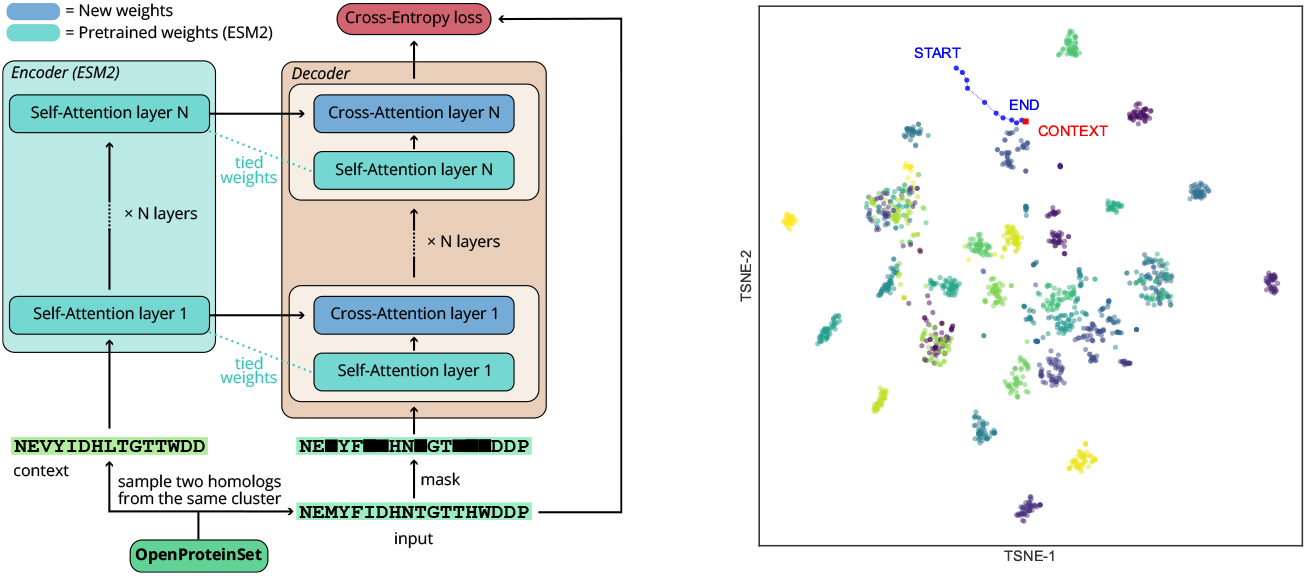
**Left:** Schematic of the RAG-ESM architecture. “Self-Attention layer” and “Cross-Attention layer” denote the concatenation of attention and feed-forward layers. **Right:** Two-dimensional t-SNE visualization of the ESM2 (650M) embeddings of protein sequences from different clusters in the test set (different colors from yellow to purple). Blue: path of a sequence generated from scratch using RAG-ESM (165M) by denoising (see Supplementary Sec. C); START: initial sequence of only **<mask>** tokens; END: sequence obtained after denoising; CONTEXT: sequence given as context.

We introduce RAG-ESM, an encoder-decoder model trained with masked language modeling on pairs of homologous protein sequences. The model takes as input both a masked sequence and a context sequence. The latter can be one of the homologs of the former, or any sequence with specific desired properties (e.g. belonging to a particular family, having a specific function, or binding to a specific domain). The context sequence is embedded using an encoder model. The resulting embeddings are provided to a decoder, in order to improve its predictions of the masked amino acids in the input sequence. We show that:

1. By conditioning on a homologous sequence, we drastically improve the performance of pretrained pLMs at predicting masked amino acids. The computational costs are minimal (50 to 120 GPU hours, depending on model size) and the number of additional parameters is small, opening new directions for more efficient and scalable future models.
2. By training the models using a discrete diffusion objective (see Section 2.2) and conditioning, during inference, on sequences with specific desired properties, we transform a pretrained single-sequence masked language model into one that can perform conditional generation.

This second contribution addresses a key limitation in the generative abilities of most single-sequence pLMs: their lack of control over generated outputs. Conditioning on sequence provides an alternative to conditioning on control tags, e.g. gene ontology terms (Nijkamp et al., 2023; Madani et al., 2023), and to multimodal pLMs using structural and functional features (Hayes et al., 2025; Wang et al., 2025). By providing a representation of a context sequence, we guide the model to sample from a specific region of sequence space, significantly reducing the dimensionality of the search space. Fig. 1 (right) shows that during the denoising process (described in Supplementary Sec. C), the model starts from a position of the embedding space corresponding to the fully masked sequence, and quickly converges to a position close to the context sequence embedding and its neighboring homologs.

Our approach reduces the need for the model to allocate parameters for memorizing protein family information (Bhattacharya et al., 2020; Vig et al., 2021; Hayes et al., 2025). Instead, this information can be recovered from the context embeddings. We find that this enables performance comparable to much larger models, with substantially fewer parameters.

Furthermore, to improve generation quality, we introduce an Error Correction strategy that allows the model to revise previously sampled residues during denoising. This leverages an MLM-like training objective (see section 2.2 and Supplementary section C), which exposes the model to corrupted inputs and trains it to detect and correct unlikely residues, even when they are not masked. In Table S6, we compare different sequence generation strategies used during denoising, showing that our proposed Error Correction method improves the quality of the output sequences.

### 2.2 Architecture and training

The RAG-ESM model builds upon a pretrained ESM2 model modified by adding a few additional layers. The architecture is shown in Fig. 1 (left). The model comprises two main modules:

1. An encoder module, which corresponds to the pretrained ESM2 model, and computes the embeddings of the (unmasked) context sequences.
2. A decoder module, built starting from the pretrained layers of the ESM2 model, adding newly initialized cross-attention layers to some of them. These layers integrate information from the context embeddings (from the encoder) and the input ones (from the decoder) into a single representation. This module takes as input the masked sequence and the embeddings of the context sequence. It provides as outputs the logits for the masked amino acids.

The weights of the ESM2 layers (including both attention and feed-forward layers) used by both encoder and decoder modules are tied, i.e. the parameters are shared between the two modules. Thus, starting with an *N*-parameter ESM2 model, and using *M* parameters in the cross-attention layers, our RAG-ESM model has *N* + *M* parameters. In practice, *M ≪ N*, as we apply cross-attention to few layers. Indeed, few layers of cross-attention suffice to transfer information between the two sequences’ embeddings, see Supplementary Sec. B. Thus motivated, we train RAG-ESM models with 12M and 165M parameters, respectively based on the 8M and 150M ESM2 models.

We train our models using the standard cross-entropy loss with a discrete diffusion objective (Sahoo et al., 2024). The masking fraction is sampled from a uniform distribution over (0, 1) to improve its generation capabilities during the denoising steps (see Supplementary Sec. A). We simultaneously fine-tune the pre-trained self-attention weights (from ESM2) and train the newly initialized cross-attention weights with different learning rates. At each training instance, a sequence is selected as input, and its closest neighbor (by Hamming distance) in the same cluster is used as context. While the final performance of the model at inference is robust to the similarity level of the context sequences used for training, using the closest neighbor during training accelerates loss convergence, see Supplementary Sec. B.

All models were trained on pairs of homologs sampled from Uniclust30 clusters (Mirdita et al., 2017) present in the filtered version of OpenProteinSet (Ahdritz et al., 2024), which comprises maximally diverse representative clusters. This dataset includes a total of ~ 200 million protein sequences from UniprotKB (The UniProt Consortium, 2021). More details are given in Supplementary Sec. A.

## 3 Results

### 3.1 Attending to close homologs drastically improves the perplexity of pLMs

Does retrieval impact the performance of pLMs? How much does this depend on the similarity between context and input sequences? We address these questions by investigating the models’ perplexity.

Fig. 2 (left) shows the effect of the distance between context and input sequences on model performance. When the two sequences are the same, the perplexity of the model is 1, i.e. the model perfectly predicts the masked amino acids by looking up those in the context sequence. Perplexity increases with the distance between the two sequences. RAG-ESM reaches perplexities similar and sometimes slightly larger than the base ESM2 model when very different sequences from the same cluster are used as context and input.

**Figure 2:**
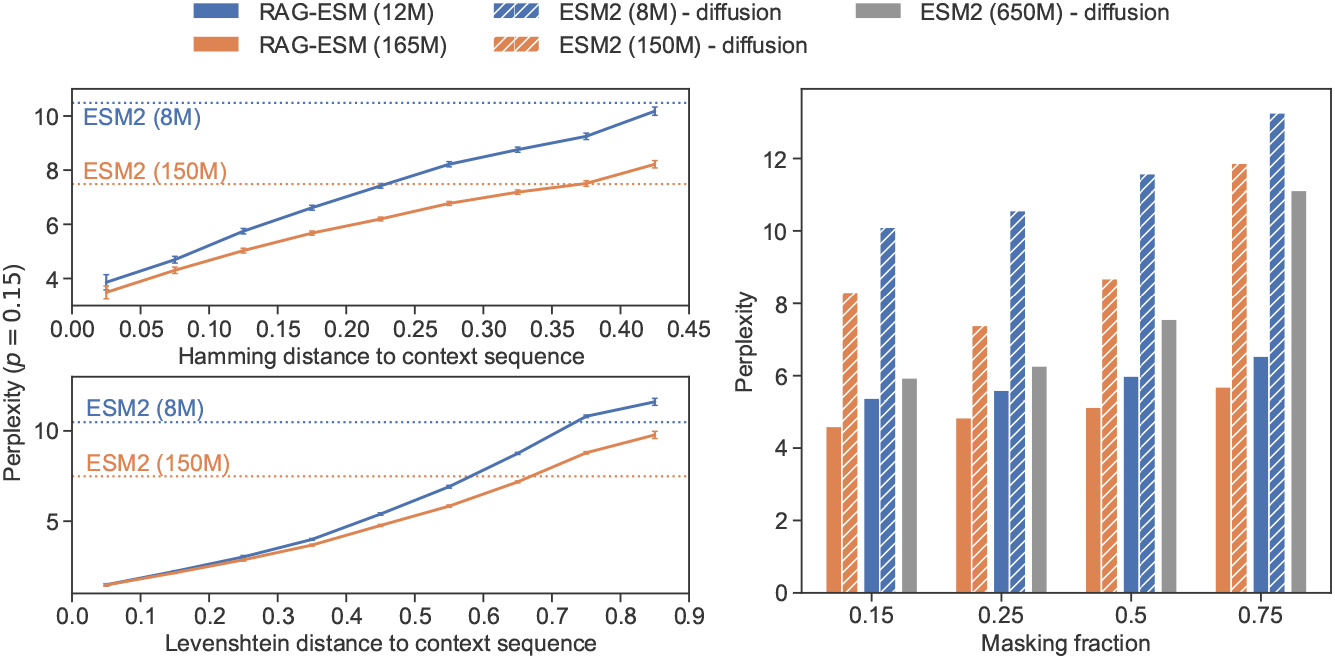
**Left:** Perplexity of the RAG-ESM models on the standard MLM task (*p* = 0.15) versus distance between input and context sequences (both from the same cluster). We measure distance using both Hamming and Levenshtein distances (resp. between sequences aligned using the pairwise Smith-Waterman algorithm, and between unaligned sequences), binned in intervals of 0.1. **Right:** Perplexity of ESM2 models and RAG-ESM models, both trained on the discrete diffusion task, for different masking fractions, when the closest homolog is provided as context to RAG-ESM (mean Hamming dist ~ 0.36). Perplexity is measured on 10,000 sequences sampled from the validation set. ESM2 (650M), fine-tuned on the discrete diffusion objective, serves as baseline (see Table S5 for a comparison with RAG-ESM models trained on other objectives).

In Fig. 2 (right), we compare RAG-ESM models with the base ESM2 models they are built on. We find that the usage of homology information leads to large performance improvements. We compare the pretrained ESM2 (8M) and ESM2 (150M), fine-tuned on the discrete diffusion task, to their RAG-ESM counterparts with 12M and 165M parameters. We obtain respectively a 48% and 43% decrease in perplexity (averaged over the different masking fractions considered in Fig. 2 (right)) with respect to the base models, when using the closest homolog as context. Performance decreases when using more distant homologs (see Table S5), but it remains at least as good as the one of ESM2 (650M). Furthermore, even the smaller RAG-ESM (12M) model reaches a lower perplexity than the much larger ESM2 (650M) model. This shows that using homologs information can help to substantially decrease the size of models, thus fostering computational efficiency.

### 3.2 Alignment capabilities naturally emerge in the cross-attention heads

How interpretable are cross-attention heads in our model? Do they learn to align input and context sequences without explicit alignment information during training? To address these questions, we examine the cross-attention coefficients calculated by RAG-ESM when feeding it pairs of homologs, without masking input sequences. Fig. 3 (top) shows that some cross-attention heads of RAG-ESM (165M) exhibit a high Pearson correlation *ρ* with pairwise sequence alignment matrices computed using a differentiable formulation proposed by (Petti et al., 2021), which was shown to converge to the Needleman-Wunsch alignment. Specifically, 7 of these heads feature *ρ >* 0.5. Note that given the sparsity of these matrices, we compute correlations *ρ* over their top 2*L* entries, where *L* is sequence length. Notably, the alignment-specialized heads appear only in the first and last cross-attention layers, while the middle layer shows little to no significant correlation. Fig. 3 (bottom) shows an alignment matrix and cross-attention matrices for two homologous sequences.

**Figure 3:**
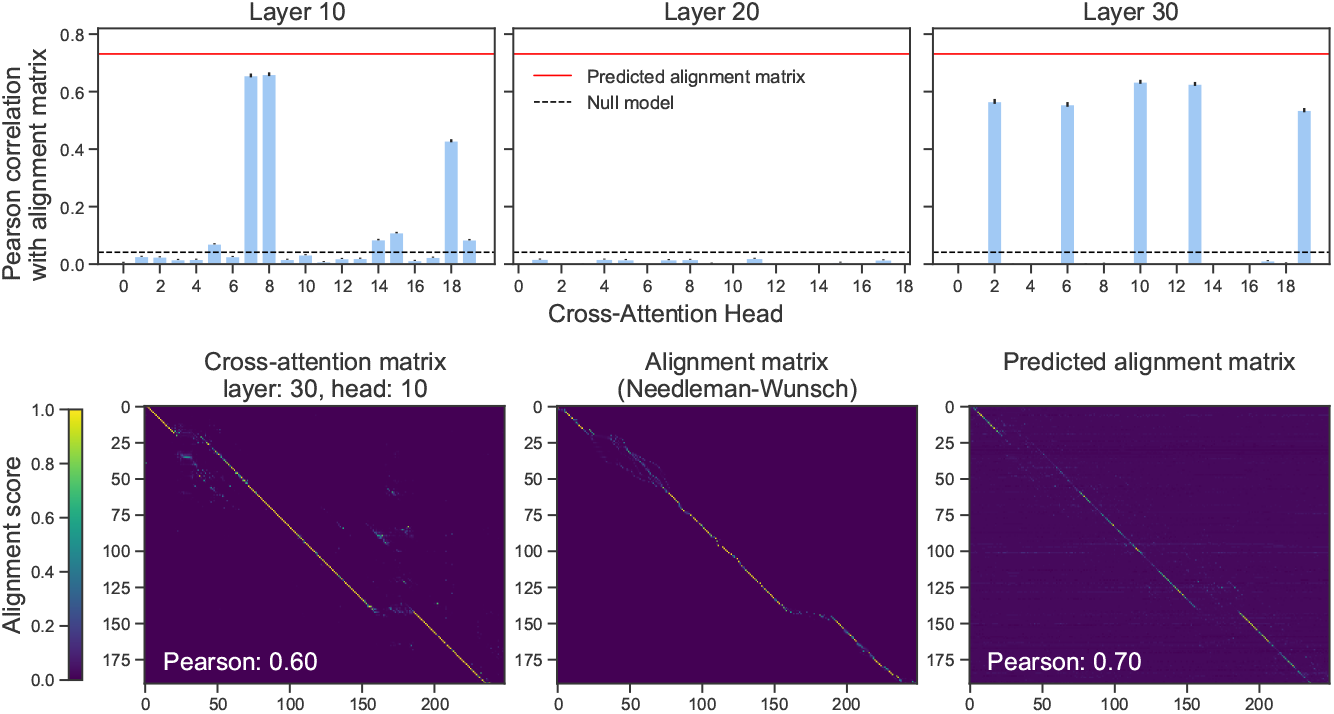
**Top:** Pearson correlation between alignment matrices obtained by aligning input and context sequences (using the Needleman-Wunsch algorithm) and each of the 60 cross-attention heads from RAG-ESM (165M) – 20 per layer across layers 10, 20, and 30. Blue and black: mean and standard error of the correlations for each head. Red line: mean Pearson correlation obtained for the logistic regression trained on 100 examples. Dashed black line: mean Pearson correlation between the alignment matrices and the same matices with shuffled entries (null model). Results are obtained by feeding one pair of homologs from each of the 1000 clusters in the test set to the RAG-ESM model, without masking the input sequences. All correlations are computed over the top 2*L* matrix entries, where *L* is sequence length. **Bottom:** Visual comparison of the Needleman-Wunsch alignment matrix (center) with one of the cross-attention heads of RAG-ESM (left) and with the alignment matrix predicted by the logistic regression (right), for a random pair of homologous sequences sampled from a cluster in the test set. Their Hamming distance is 0.73.

To further assess the ability of attention heads at capturing alignment matrices, we trained a logistic regression model (with predictions ranging continuously between 0 and 1) on the 60 cross-attention heads to predict the alignment matrices. Using 100 samples from the test set for training, and evaluating on the remaining 900 samples, we obtained an average Pearson correlation of *ρ* = 0.73 between the alignment matrices predicted by the logistic model and the original ones, see Fig. 3 (top). This result is reminiscent of Rao et al. (2021), where logistic regression on row attention matrices was shown to capture contact maps, and of Lupo et al. (2022), where logistic regression on column attention matrices was found to predict Hamming distances between sequences.

These results suggest that the cross-attention between the input and context sequences enables the model to implicitly perform sequence alignment, despite not being trained on explicitly aligned sequences. In other words, the training objective of RAG-ESM encourages the extraction of informative signals from homologous context sequences. This leads to the emergence of specialized cross-attention heads that tend to effectively align the two sequences.

### 3.3 Diffusion-based denoising enables conditional protein sequence generation

RAG-ESM can generate protein sequences conditioned on homologous context via a diffusion-based denoising process that progressively reveals masked parts of the input sequences (see Supplementary Sec.C for details). How does retrieval influence generative performance? To answer this, we compare sequences generated by RAG-ESM (165M) with those produced by a diffusion fine-tuned ESM2 (650M) baseline, which does not directly use homolog information, and with natural sequences from the test set. The natural sequences consist of (i) homologous ones sampled from the same clusters as the generated sequences, and (ii) non-homologous ones sampled from random clusters.

To assess the quality of the generated sequences, we employ several established metrics. Sequence confidence is measured via the pseudo-perplexity of the non-fine-tuned ESM2 (650M), while structural confidence is quantified by the ESMFold pLDDT (Lin et al., 2023) and the ProteinMPNN self-consistency perplexity (scPerplexity, computed from ESMFold-predicted structures, following Alamdari et al. (2023)). In addition, structural similarity between the predicted structures of generated and context sequences is evaluated using RMSD and TMScore (after structural alignment) (Zhang & Skolnick, 2004), and homology is assessed using HMMER scores (Eddy, 2020) obtained from a hidden Markov model trained on the cluster’s MSA. Finally, we use the Hamming distance to quantify how much each generated sequence diverges from the corresponding context sequence.

The results in Table 1 indicate that RAG-ESM (165M) significantly outperforms the larger ESM2 baseline for conditional sequence generation. This finding confirms the benefit of incorporating homologs to improve model performance during inference. Notably, RAG-ESM-generated sequences have scores closer to those of natural sequences, and sometimes exceed them. Specifically, the HMMER scores of RAG-ESM-generated sequences closely approximate those of natural homologs, highlighting the model’s ability to preserve key characteristics of protein families. Moreover, the Hamming distances between generated sequences and their corresponding context sequences are comparable to those observed between natural homologs, indicating that our method produces sequences that appropriately differ from the provided context. In terms of structural plausibility, the sequences generated by RAG-ESM feature ESMFold pLDDT scores and ProteinMPNN self-consistency perplexities that are similar to those of natural sequences, and superior to those generated by the larger ESM2 baseline. Although the ESM2 pseudo-perplexity for RAG-ESM is marginally less favorable than that of natural sequences, it still outperforms the ESM2 baseline. Additionally, RAG-ESM produces sequences that are structurally more similar to their context than ESM2, as evidenced by significantly lower RMSD and higher TMScore values compared to both natural homologs and ESM2-generated sequences. Thus, the conditional generation strategy yields enhanced structural performance.

**Table 1:** Quality scores for generated and natural sequences from 1000 different clusters in the test set. We report the median and standard deviation of each distribution. Both generated and natural sequences are filtered by computing the ESM2 (650M) pseudo-perplexity and retaining only those with perplexity *<* 8 (approximately 650 sequences out of 1000, both for natural and generated sequences). HMMER scores are normalized by dividing each of them by the score of the corresponding context sequence. For natural sequences, Hamming distances to the context are computed for randomly sampled sequences, either from the same cluster as the context (“Homolog”) or from a random different cluster (“Random”). *↑* (resp. *↓*) indicates that higher (resp. lower) scores are better.

| Score | Generated |  | Natural |  |
| --- | --- | --- | --- | --- |
|  | RAG-ESM (165M) | ESM2 (650M) | Homolog | Random |
| ESMFold pLDDT (↑) | $0.76 \pm 0.12$ | $0.52 \pm 0.16$ | $0.81 \pm 0.15$ | |
| scPerplexity (↓) | $2.58 \pm 0.44$ | $2.96 \pm 0.68$ | $2.52 \pm 0.43$ | |
| ESM2 (650M) perplexity (↓) | $6.69 \pm 2.66$ | $7.16 \pm 2.47$ | $6.06 \pm 3.03$ | |
| RMSD with context (↓) | $4.49 \pm 5.41$ | $13.13 \pm 7.76$ | $9.68 \pm 6.0$ | $11.85 \pm 4.5$ |
| TMScore with context (↑) | $0.49 \pm 0.26$ | $0.14 \pm 0.05$ | $0.19 \pm 0.15$ | $0.15 \pm 0.05$ |
| HMMER (↑) | $0.96 \pm 0.76$ | $-0.01 \pm 0.25$ | $0.92 \pm 0.62$ | $-0.01 \pm 0.1$ |
| Hamming to context ($-$) | $0.50 \pm 0.21$ | $0.74 \pm 0.04$ | $0.50 \pm 0.16$ | $0.77 \pm 0.05$ |

Table 2 further compares the performance of RAG-ESM for conditional generation with other protein language models. ProtMamba (Sgarbossa et al., 2024), currently state-of-the-art for conditional generation, is a Mamba-based autoregressive model that leverages evolutionary information during inference by concatenating multiple unaligned homologous sequences as context. EvoDiff-MSA is an MSA-based diffusion protein language model whose capabilities have been experimentally validated (Alamdari et al., 2023). MSA Transformer is the pretrained model used as the starting point for EvoDiff-MSA and was trained with a masked language modeling objective (Rao et al., 2021). Finally, we consider Potts models trained on the MSAs of specific families, an experimentally validated generative model inspired by statistical physics (Russ et al., 2020a). We compare RAG-ESM with these baseline models using the performance reported in (Alamdari et al., 2023) and (Sgarbossa et al., 2024). Strikingly, RAG-ESM achieves the highest median ESMFold pLDDT and the lowest ProteinMPNN scPerplexity among the generative models considered. Thus, RAG-ESM performs as well as the state-of-the-art method ProtMamba, even if it conditions on single sequences instead of larger sets of homologs from the same cluster. Furthermore, RAG-ESM exceeds the performance of other existing models for homolog-conditioned generation.

**Table 2:** Performance of RAG-ESM and other models for homolog-conditioned generation. We report the median and standard deviation of each distribution. Two structural scores are presented: the pLDDT from ESMFold (Lin et al., 2023) and the scPerplexity from ProteinMPNN (Dauparas et al., 2022), evaluated on a set of 250 protein sequences generated using each of the models, each from a distinct cluster in the test set. For other models than RAG-ESM, results were retrieved from the Zenodo archive associated with the EvoDiff paper (Alamdari et al., 2023) and the ProtMamba paper (Sgarbossa et al., 2024). *↑* (resp. *↓*) indicates that higher (resp. lower) scores are better.

| Model | RAG-ESM | ProtMamba | EvoDiff-MSA | MSA Trans. | Potts | Natural |
| --- | --- | --- | --- | --- | --- | --- |
| pLDDT (↑) | **$0.76 \pm 0.12$** | $0.75 \pm 0.13$ | $0.60 \pm 0.16$ | $0.54 \pm 0.18$ | $0.56 \pm 0.14$ | $0.81 \pm 0.15$ |
| scPerplexity (↓) | **$2.58 \pm 0.44$** | $2.63 \pm 0.45$ | $3.17 \pm 0.58$ | $3.37 \pm 0.64$ | $3.17 \pm 0.51$ | $2.52 \pm 0.43$ |

Together, these results, along with the high cosine similarity observed between ESM2 (650M) embeddings of generated and context sequences (Fig. S1), demonstrate that the diffusion-based denoising strategy of RAG-ESM effectively leverages homologous information for conditional generation. The approach yields protein sequences that are both evolutionarily and structurally plausible, performing similarly to natural sequences across the evaluated metrics. This positions RAG-ESM as a promising alternative to existing generation methods, particularly in applications where preserving context-specific structural and evolutionary characteristics is critical.

### 3.4 RAG-ESM reaches state-of-the-art performance for motif scaffolding

Motivated by the promising results in conditional sequence generation, we assess the generative performance of RAG-ESM (165M) for the motif-scaffolding task introduced by Watson et al. (2023). The goal is to design protein scaffolds that accommodate a fixed functional motif, such as a binding or function-determining region, by generating the surrounding sequence that supports its structural integrity. Fig. 4 shows examples of motifs that were successfully scaffolded via denoising with RAG-ESM.

**Figure 4:**
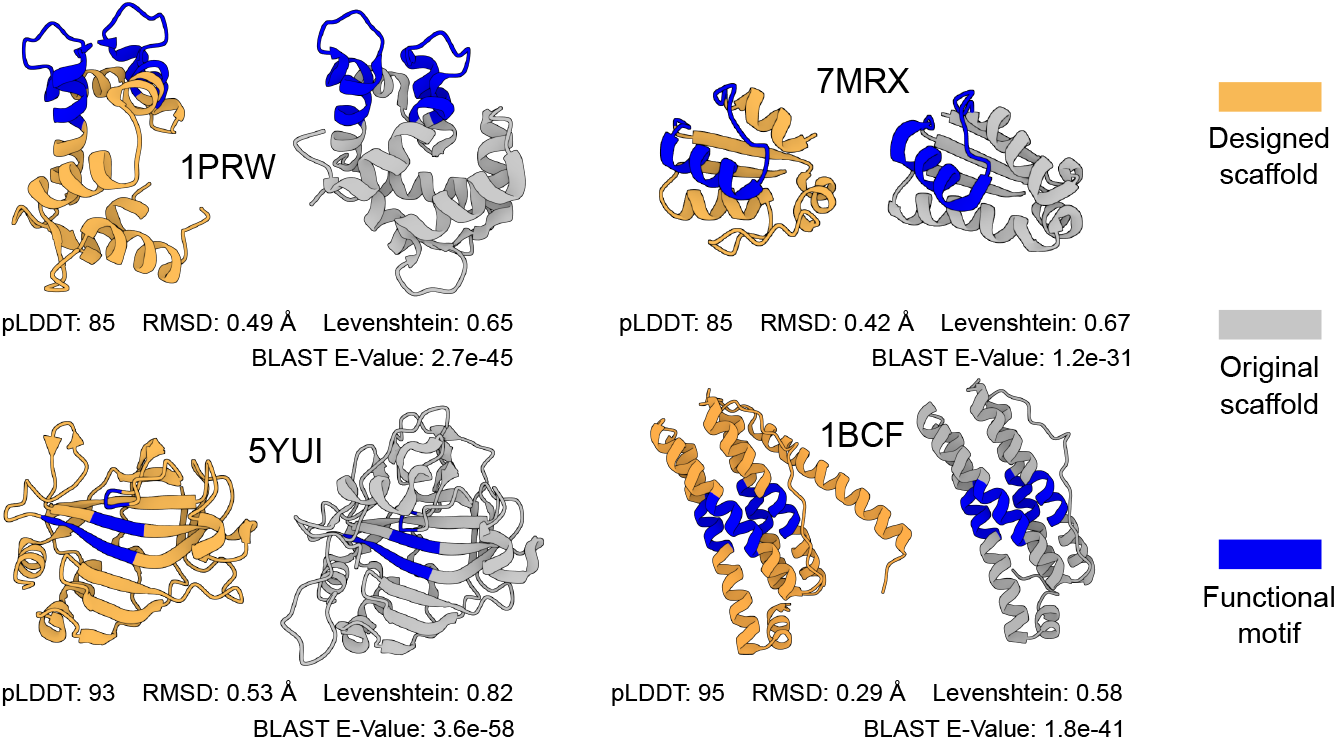
Examples of successfully scaffolded motifs. The structure of each motif is highlighted in blue, while gray denotes the reference structures from the PDB, and orange denotes the ESMFold-predicted structures for the designed scaffolds. For each design, we report the ESMFold pLDDT (higher is better), the RMSD – restricted to the motif region – between generated and reference structure (lower is better), the Levenshtein distance between generated and reference sequence (higher is better), and the BLAST E-Value score when searching the UniProtKB database (The UniProt Consortium, 2021). Note that the similarity between the designed sequences and the closest natural sequence from UniprotKB, identified by the BLAST search, are the following: 1BCF: 59.5%; 7MRX: 68.4%; 5YUI: 56.7%; 1PRW:62.8%.

Here, we follow the procedure proposed in (Alamdari et al., 2023), with one modification: we use ESMFold (Lin et al., 2023) as the structure prediction model instead of OmegaFold (Wu et al., 2022), motivated by the higher accuracy of the former. We sample the new scaffolds by conditioning RAG-ESM on the original sequence of each motif and using a temperature of *T* = 0.7.

How does RAG-ESM compare to other models for motif scaffolding? Previous approaches include sequence-based methods like DPLM (650M) (Wang et al., 2025), the alignment-based EvoDiff-MSA (Alamdari et al., 2023), structure-based models such as RFDiffusion (Watson et al., 2023), and large multimodal models incorporating structural information like ESM3 (1.4B) (Hayes et al., 2025). In Table 3, we compare the success rates of RAG-ESM (165M) to these methods for the motif-scaffolding task. Our results show that RAG-ESM outperforms the larger DPLM (650M) model, as well as the MSA-based EvoDiff-MSA. Although the structure-based model RFDiffusion and the multimodal model ESM3 remain superior on some motifs, they are outperformed by RAG-ESM on others. Thus, we envision that RAG-ESM could complement structure-based approaches.

**Table 3:**
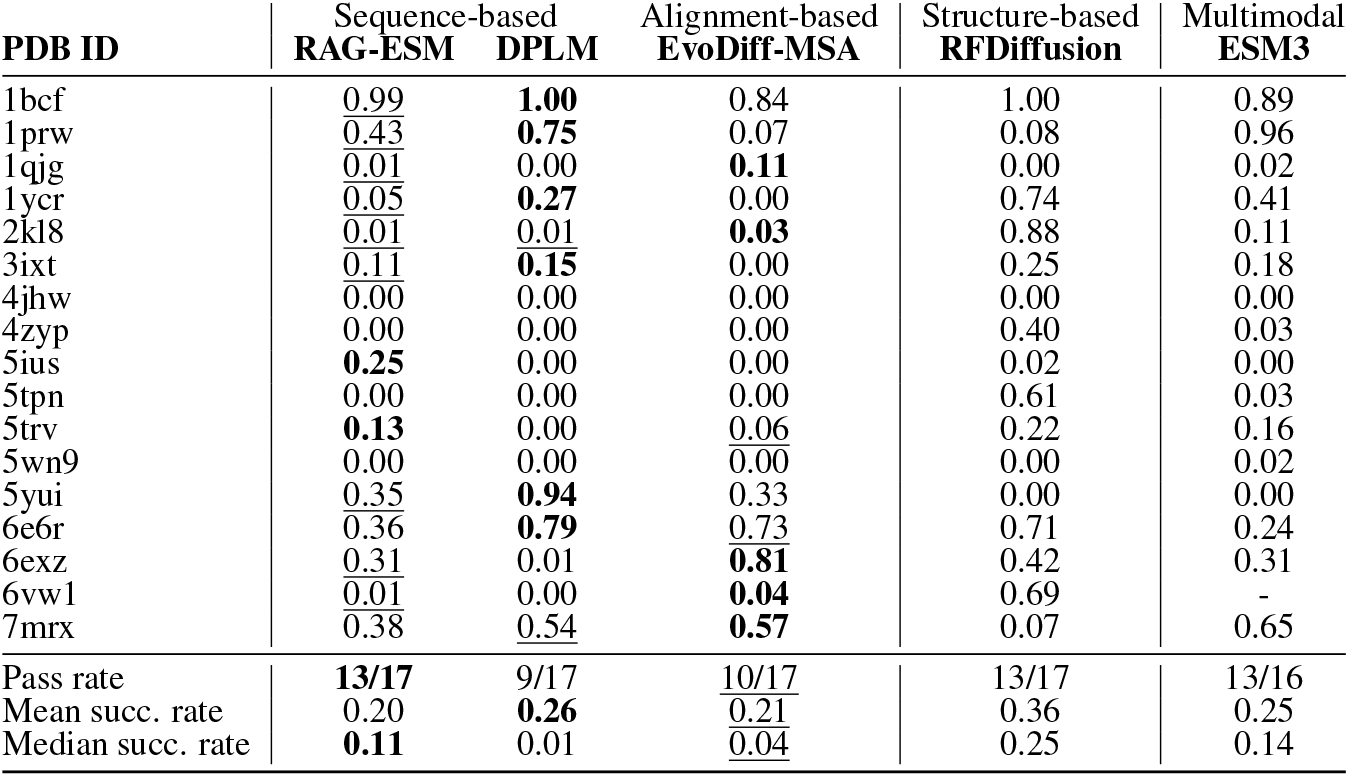
Performance comparison of different models on the motif-scaffolding task. For each of the 17 motifs considered in Watson et al. (2023), labeled by the associated PDB entry, we report the success rate (out of 100 attempts for each target) of scaffold generation for five models: RAG-ESM (165M), DPLM (650M), EvoDiff-MSA, RFDiffusion, and ESM3 (1.4B). Scaffold sequences are generated with lengths sampled uniformly from [50, 100] plus the motif length, and subsequently folded using ESMFold, similarly to (Alamdari et al., 2023). A design is considered successful if its folded structure exhibits a pLDDT score greater than 0.70 and an RMSD to the motif (in the reference structure) smaller than 1 Angstrom. We also report the overall pass rate (i.e., the number of tasks with at least one successful design), as well as the mean and median success rates. Both are reported because the distribution of success rates across motifs is highly skewed. Models with the best performance are in **bold**, and those with the second-best one are <u>underlined</u>. For ESM3, we show values reported in (Wang et al., 2024), thus omitting 6vw1.

How robust are the comparisons between models presented here? One challenge for detailed comparison between sequence-based and structure-based models is that structure-based models like RFDiffusion follow a different pipeline for motif scaffolding. They usually generate a structural scaffold that is subsequently inverse-folded using ProteinMPNN. We nevertheless included these comparisons here for completeness. Besides, there appear to be some differences in the reported performance of DPLM for motif-scaffolding tasks between (Wang et al., 2025) and the more recent study (Wang et al., 2024). For completeness, in Table S7, we compare our results with the updated DPLM performance (Wang et al., 2024). That comparison is even more favorable to RAG-ESM than Table 3. Finally, in Table S8, we present a similar comparison as in Table 3, but using OmegaFold instead of ESMFold as the structure prediction method. This yields higher success rates due to OmegaFold’s higher pLDDT scores, but does not modify our conclusions on the comparisons between models. Thus, our conclusions are robust, and show that RAG-ESM is very promising for motif scaffolding.

## 4 Discussion

In this work, we introduced RAG-ESM, a sequence retrieval-augmented framework that builds upon a pretrained protein language model trained on an unstructured ensemble of protein sequences, specifically ESM2. Specifically, we augmented ESM2 with a small number of cross-attention layers. RAG-ESM takes as input a masked sequence and as context a homologous sequence, and computes cross-attention between them. We showed that such conditioning on homologous sequences dramatically reduces perplexity, and enables relatively small models to perform on par with much larger ones. We also found that some cross-attention heads of RAG-ESM spontaneously learn to align input and context sequences. Thus, RAG-ESM efficiently learns sequence homology.

By integrating a discrete diffusion objective, we enabled RAG-ESM to perform conditional protein sequence generation. We showed that RAG-ESM not only achieves state-of-the-art performance in generating novel sequences, but also excels in the motif-scaffolding task among sequence-based models. In fact, it outperforms models that either use significantly more parameters (Wang et al., 2025) or rely on additional information, such as MSA-based models (Alamdari et al., 2023). An asset of RAG-ESM is that it strongly benefits from using just one homologous sequence as context, while other models that perform homology-conditioned generation require MSAs (Rao et al., 2021) or large collections of homologs (Truong Jr & Bepler, 2024; Sgarbossa et al., 2024), and models that condition on control tags need annotated data for training (Madani et al., 2020; 2023; Munsamy et al., 2022). This makes RAG-ESM more flexible, enhancing its applicability e.g. to small protein families and poorly annotated data.

Our work opens several promising future directions. For instance, incorporating additional sources of biological information, such as structural data or functional annotations, could further improve the controllability and accuracy of generated sequences (Su et al., 2023; Heinzinger et al., 2024; Hayes et al., 2025; Michalewicz et al., 2025). Moreover, exploring more advanced denoising algorithms might enhance the generative capabilities of the model even further (Sahoo et al., 2024; Wang et al., 2025). Besides, the emergent alignment ability of RAG-ESM suggests its potential for developing new methods for unsupervised protein alignment and evolutionary analysis. Finally, RAG-ESM establishes a scalable, efficient, and versatile protein sequence generation method conditioned on homologous sequences. It thus opens new avenues for controlled protein engineering.

## Data availability statement

A Python implementation of RAG-ESM is freely available in our GitHub repository: https://github.com/Bitbol-Lab/rag-esm

## Acknowledgments

This research was partly funded by the European Research Council (ERC) under the European Union’s Horizon 2020 research and innovation programme (grant agreement No. 851173, to A.-F. B.).

## Supplementary information

### A. Training details

The models are trained using the standard cross-entropy loss on the masked amino acids of the input sequence. We train them using two different training regimes (compared in Table S5):

1. Standard Masked Language Modeling (MLM): masking fraction *p* = 0.15 (Devlin et al., 2019).
2. Discrete diffusion objective: we experiment with two different masking techniques, the first is the standard discrete diffusion objective where the masking fraction is sampled from a uniform distribution over (0, 1), in the second we sample the masking fraction 80% of the time from a *β*(3, 9) distribution and 20% of the time from a uniform distribution over (0, 1). This approach, adapted from (Hayes et al., 2025), aims to balance representation and generation capabilities. It allows the model to observe masking fractions across (0, 1), with an average 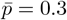. Both these objectives improve the effectiveness for iterative denoising during sequence generation with respect to standard MLM (Alamdari et al., 2023; Wang et al., 2025).

We employ the AdamW optimizer with the following parameters: weight decay *w* = 0.1 and (*β*_1_, *β*_2_) = (0.9, 0.98). Learning rates are set to 1 *×* 10^−4^ for pretrained ESM2 self-attention weights and 1 *×* 10^−3^ for newly initialized cross-attention weights. Additional information on the ablation studies performed to choose the training parameters and configuration is provided in Section B. To optimize memory usage, we use mixed precision with `bfloat16`. Dropout was not used because our ablation studies showed no benefit for loss convergence or overfitting.

All models are trained on single NVIDIA RTX 6000 GPUs with an effective batch size of 1024, obtained via gradient accumulation, for 21k steps (in practice, in a full training run, a model sees 100 homologs per protein sequence cluster, see Sec. 2.2). Input and context sequences are cropped to a maximum length of 512 tokens. Since ESM2 uses rotary positional embeddings, the model can handle significantly longer sequences during inference (Lin et al., 2023). We checked that lengths up to 2048 tokens could be used without performance degradation. Finally, to reduce training time, we replace the slower ESM2 attention implementation with PyTorch’s scaled dot product attention (i.e. FlashAttention).

#### Dataset

All models are trained on OpenProteinSet (Ahdritz et al., 2024), a dataset comprising 16 million MSAs, each representing a sequence cluster from the clustered sequence database Uniclust30 (Mirdita et al., 2017). This dataset, curated for training OpenFold (Ahdritz et al., 2022), was filtered to include only maximally diverse representative MSA clusters. Redundant clusters, whose representative sequences appeared in other clusters’ MSAs, were iteratively removed (Ahdritz et al., 2024). As a result, each representative sequence is unique to its cluster, as detailed in Ahdritz et al. (2024). The filtered dataset includes 268,000 clusters, totaling ~ 200 million protein sequences from UniprotKB (The UniProt Consortium, 2021). Validation and testing sets are each created by holding out 1,000 randomly selected clusters from the training set. Importantly, the filtering minimizes overlap between clusters in the training, validation, and test sets. By focusing on MSAs of maximal diversity and ensuring that reference sequences are unique to their clusters, this dataset ensures strong partitioning into diverse clusters of homologs.

During training, a sequence is randomly sampled from a cluster to be the model’s input, and its closest neighbor (i.e. the sequence with the lowest Hamming distance within the same cluster) is used as the context sequence. Ablation studies on different training modalities (see Section B) show that the model’s final performance remains similar regardless of how close the context sequences are to the input ones during the training phase. In other words, using distant homologs has the same effect on the final performance as using the closest ones. Our motivation for choosing the closest neighbors as context is that they lead to a faster loss convergence.

### B Ablations

We tested different training configurations, using the smallest ESM2 model (8M parameters) unless specified otherwise (e.g. in Tables S3 and S5 we also show ablations for ESM2 (150M)). In all the tables, we denote in *italics* the performance of pretrained ESM2 models, which serve as a baseline, to distinguish them from the ablations that we performed. We also highlight in **bold** the models with best performance and <u>underline</u> those with second best performance for each task. Finally, we denote by “Perplexity closest” and “Perplexity random” the perplexities of the model in the MLM task (with *p*_*mask*_ = 0.15) when using as context sequence either the closest homolog or a random homolog sampled from the same cluster as the input.

The ablation studies in Table S1 show that the model’s final performance remains similar regardless of how close the context sequences are to the input ones during the training phase. Different rows in the table correspond to different types of homologs used as context during training, while different columns correspond to different evaluation sets (see previous paragraph).

**Table S1:** Using different types of context sequences as training set.

| Context sequence sampling (training) | Perplexity closest | Perplexity random |
| --- | --- | --- |
| *ESM2 (8M) – No context* | *10.48* | *10.48* |
| Closest neighbor | **5.54** | 8.55 |
| Top-10 closest neighbors | <u>5.58</u> | **8.51** |
| Random homologs | 6.20 | 8.65 |

The ablations in Table S2 show that the optimal training configuration is the one where self-attention (pretrained), cross-attention (randomly initialized) and encoder (pretrained) are all trained together. Since tying the weights of the self-attention layers of encoder and decoder largely decreases the number of effective parameters with minimal effects on the performance, we decided to train the final models using the tied configuration.

**Table S2:** Different training configurations and trained parameters. Same conventions as in Table S1. C-A: cross-attention; S-A: self-attention.

| Training configuration | Trained parameters | Perplexity closest | Perplexity random |
| --- | --- | --- | --- |
| *ESM2 (650M)* | - | *6.13* | *6.13* |
| *ESM2 (8M) – pretrained* | - | *10.48* | *10.48* |
| Train only C-A | 7.4M | 5.54 | 8.55 |
| Train S-A & C-A | 7.8M + 7.2M | 5.42 | 8.45 |
| Train S-A, C-A & Encoder | 7.8M + 7.2M + 7.8M | **5.37** | **8.35** |
| Train S-A, C-A & Encoder (tied) | 7.8M + 7.2M | <u>5.41</u> | <u>8.40</u> |

In Table S3, we compare models trained using different numbers of cross-attention layers, both in the 8M and in the 150M parameters models. We find that it is not necessary to use cross-attention after each self-attention layer. In fact, decreasing the number of cross-attention layers brings no loss in performance of the models, although convergence is slower. We also study the case in which cross-attention is applied only to the last layer of the decoder (similarly to Zheng et al. (2023)), but this leads to worse performance than our other methods, although it remains much better that the base ESM2 model. Based on the results in Table S3, we decided to train the final RAG-ESM model based on ESM2 (8M) using cross-attention every other layer, and the RAG-ESM model based on ESM2 (150M) using cross-attention every 10 layers (i.e. at layers 10, 20 and 30).

**Table S3:**
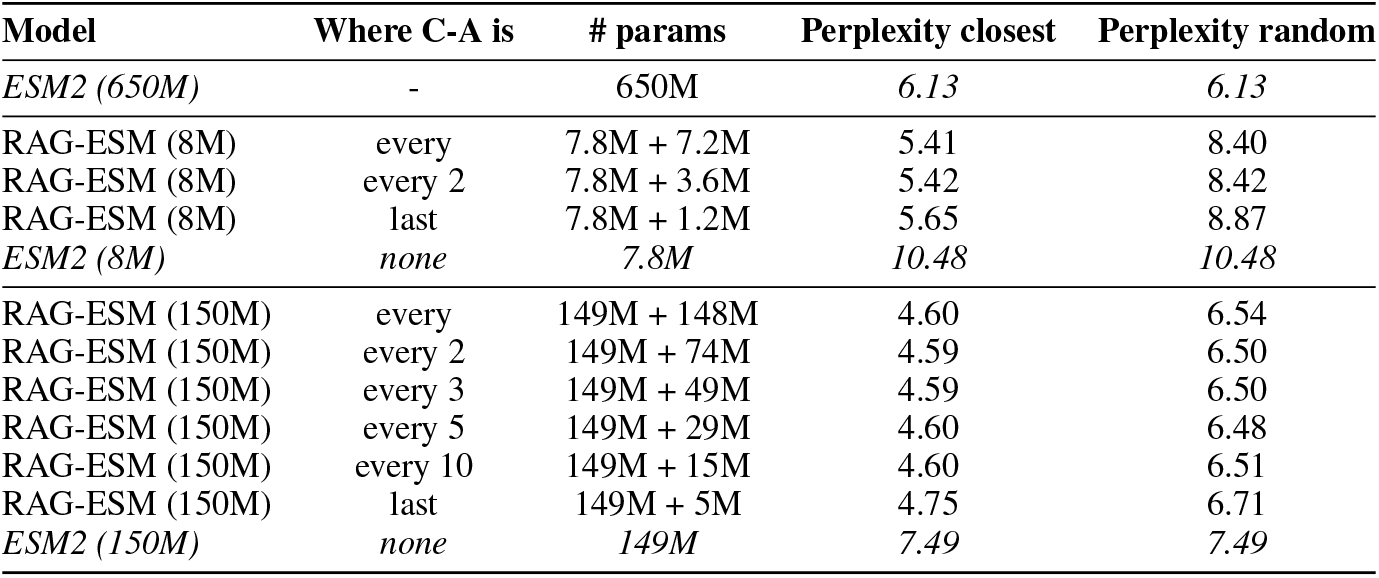
Different ways of interleaving cross-attention and self-attention.

Motivated by the ablations in table S4, we chose the learning rates to be 1 *×* 10^−4^ for the pretrained self-attention layers and 1 *×* 10^−3^ for the newly initialized cross-attention layers. Recall that ESM2 models were all pretrained using a peak learning rate of 4 *×* 10^−4^ (see (Lin et al., 2023)).

**Table S4:**
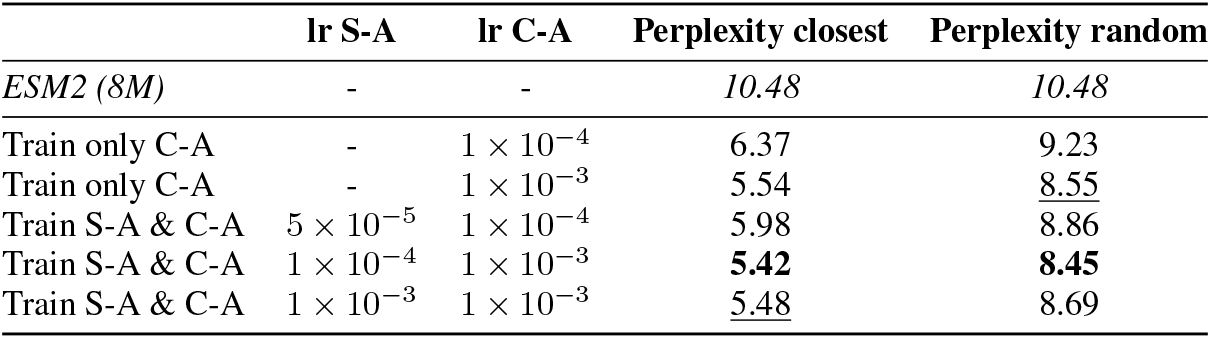
Different learning rates (lr).

|  | lr S-A | lr C-A | Perplexity closest | Perplexity random |
| --- | --- | --- | --- | --- |
| *ESM2 (8M)* | - | - | *10.48* | *10.48* |
| Train only C-A | - | $1 \times 10^{-4}$ | 6.37 | 9.23 |
| Train only C-A | - | $1 \times 10^{-3}$ | 5.54 | <u>8.55</u> |
| Train S-A & C-A | $5 \times 10^{-5}$ | $1 \times 10^{-4}$ | 5.98 | 8.86 |
| Train S-A & C-A | $1 \times 10^{-4}$ | $1 \times 10^{-3}$ | **5.42** | **8.45** |
| Train S-A & C-A | $1 \times 10^{-3}$ | $1 \times 10^{-3}$ | <u>5.48</u> | 8.69 |

In Table S5, we compare different training objectives for RAG-ESM models, namely the standard masked language modeling (MLM) objective with masking fraction *p* = 0.15, and two discrete diffusion objectives, one where the masking probability is uniformly sampled from a uniform distribution over (0, 1) named “Diffusion (uniform)”, and one where it is sampled 80% of the time from a *β*(3, 9) distribution and 20% of the time from a uniform distribution over (0, 1) as in (Hayes et al., 2025), named “Diffusion (ESM3-style)”. We observe that the training objective that has the better performance on a wide range of masking fractions are the discrete diffusion ones, with little difference between the two ways of choosing masking probability. We decide to train models using the “Diffusion (uniform)” one since it is the standard in literature. Furthermore, this objective is preferred to MLM because it allows the models to be used for the generative task via iterative denoising. Table S5 further shows a comparison between the ESM2 models, both the pretrained ones and the ones fine-tuned on the diffusion task, and their RAG-ESM counterparts trained on the same objectives. This comparison shows that even when ESM2 models are fine-tuned for the diffusion task, leveraging homology information has a strong effect on performance, and gives 43% to 48% improvements with respect to ESM2 models.

**Table S5:** Different training objectives.

| Masking fraction: | Perplexity closest neighbors |  |  |  | Perplexity random homologs |  |  |  |
| --- | --- | --- | --- | --- | --- | --- | --- | --- |
|  | 0.15 | 0.25 | 0.5 | 0.75 | 0.15 | 0.25 | 0.5 | 0.75 |
| ESM2 (8M) - MLM | 10.48 | 11.21 | 14.01 | 20.42 | 10.48 | 11.21 | 14.01 | 20.42 |
| ESM2 (150M) - MLM | 7.57 | 8.16 | 11.28 | 18.82 | 7.57 | 8.16 | 11.28 | 18.82 |
| ESM2 (650M) - MLM | 6.13 | 6.66 | 9.45 | 36.68 | 6.13 | 6.66 | 9.45 | 36.68 |
| ESM2 (8M) - diffusion | 10.10 | 10.56 | 11.58 | 13.26 | 10.10 | 10.56 | 11.58 | 13.26 |
| ESM2 (150M) - diffusion | 8.30 | 7.39 | 8.68 | 11.87 | 8.30 | 7.39 | 8.68 | 11.87 |
| ESM2 (650M) - diffusion | 5.94 | 6.27 | 7.56 | 11.12 | 5.94 | 6.27 | 7.56 | 11.12 |
| **RAG-ESM (12M)** |  |  |  |  |  |  |  |  |
| MLM ($p = 0.15$) | 5.41 | <u>5.65</u> | 6.28 | 9.00 | **8.40** | <u>8.60</u> | 9.96 | 12.65 |
| Diffusion (ESM3-style) | <u>5.38</u> | **5.60** | <u>5.99</u> | <u>6.54</u> | <u>8.40</u> | **8.54** | **9.49** | <u>10.77</u> |
| Diffusion (uniform) | **5.31** | 5.70 | **5.92** | **6.49** | 8.54 | 8.93 | <u>9.81</u> | **10.67** |
| **RAG-ESM (165M)** |  |  |  |  |  |  |  |  |
| MLM ($p = 0.15$) | 4.81 | 5.24 | 6.83 | 12.90 | 6.61 | 6.79 | 8.89 | 14.06 |
| Diffusion (ESM3-style) | 4.60 | 4.84 | 5.13 | 5.69 | 6.51 | 6.60 | 7.61 | 9.42 |

Finally, we evaluate whether the model is biased towards residues that are conserved with respect to mutated ones. To this end, we compare the performance of the model (via the perplexity), under the same conditions as in the other ablations, when masking two different types of residues in the input sequence:

1. Residues that are identical to those in the context sequence, as determined from a Smith-Waterman alignment of the two sequences.
2. Residues that differ between these two sequences.

We obtain a perplexity of 4.68 in the first case and 4.61 in the second. Hence, the performance of the model is not better for conserved residues. This shows that the model’s improved performance, with respect to the baseline ESM2 model, is not due to copying residues from the context sequence.

### C Sequence generation via iterative denoising

The RAG-ESM models, trained with a discrete diffusion objective (see Sec. 2.2 for details), can be used to generate novel sequences conditioned on the context. Building on prior work (Zheng et al., 2023; Alamdari et al., 2023; Wang et al., 2025), we developed a simple denoising algorithm for sampling sequences from these models. The process begins by selecting the total number *T* of timesteps and preparing the input sequence, which may be fully or partially masked. Let the *i*-th token of the masked sequence of length *L* be *x*_*i*_ for *i* ∈ {1, …, *L*}. Let *M* denote the total number of masked tokens in the sequence and *f*_*θ*_ denote the model. At each timestep *t* ∈ {0, …, *T*}, the sequence is denoised as follows:

1. Logit prediction: The masked sequence 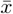 is passed to the model *f*_*θ*_, yielding logits 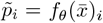 for each position *i*. The logits are then transformed into a categorical distribution of amino acid probabilities: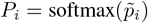.
2. Token replacement: A fraction *M/T* of the masked tokens is replaced with tokens sampled from the predicted distribution, *x*_*i*_ *∼ P*_*i*_. The tokens to replace can be selected randomly or prioritized based on the distributions *P*_*i*_ with the lowest entropy (i.e., the most confident predictions).
3. Error correction (optional): For all non-mask tokens *x*_*i*_, if the amino acid corresponding to the maximum probability in *P*_*i*_ differs from *x*_*i*_, the token is updated as *x*_*i*_ = argmax(*P*_*i*_). This step allows the model to revise previous predictions that may not be compatible anymore with the evolving sequence.

The model’s ability to perform error correction (step 3) is allowed by its training strategy, which uses the standard masking approach of the original BERT model (Devlin et al., 2019), albeit with a different way of selecting masking probabilities (see A). During training, masked tokens are replaced 80% of the time by **<mask>**, 10% by the original token and 10% by a random token. This strategy helps the model to identify incorrect tokens and propose suitable replacements, even for unmasked inputs, during the denoising steps.

Using this denoising approach, RAG-ESM-generated sequences consistently converge to regions of the embedding space close to the context sequence, allowing for precise conditional sequence generation. In Fig. S1, we show the cosine similarity between the embeddings of generated sequences and context sequences at every denoising step. In the vast majority of cases, the generated sequences converge to the right region of the sequence space (i.e., they feature a cosine similarity with the corresponding context sequence that increases towards 1). See also Fig. 1 (right) for a visualization of the denoising process.

**Figure S1:**
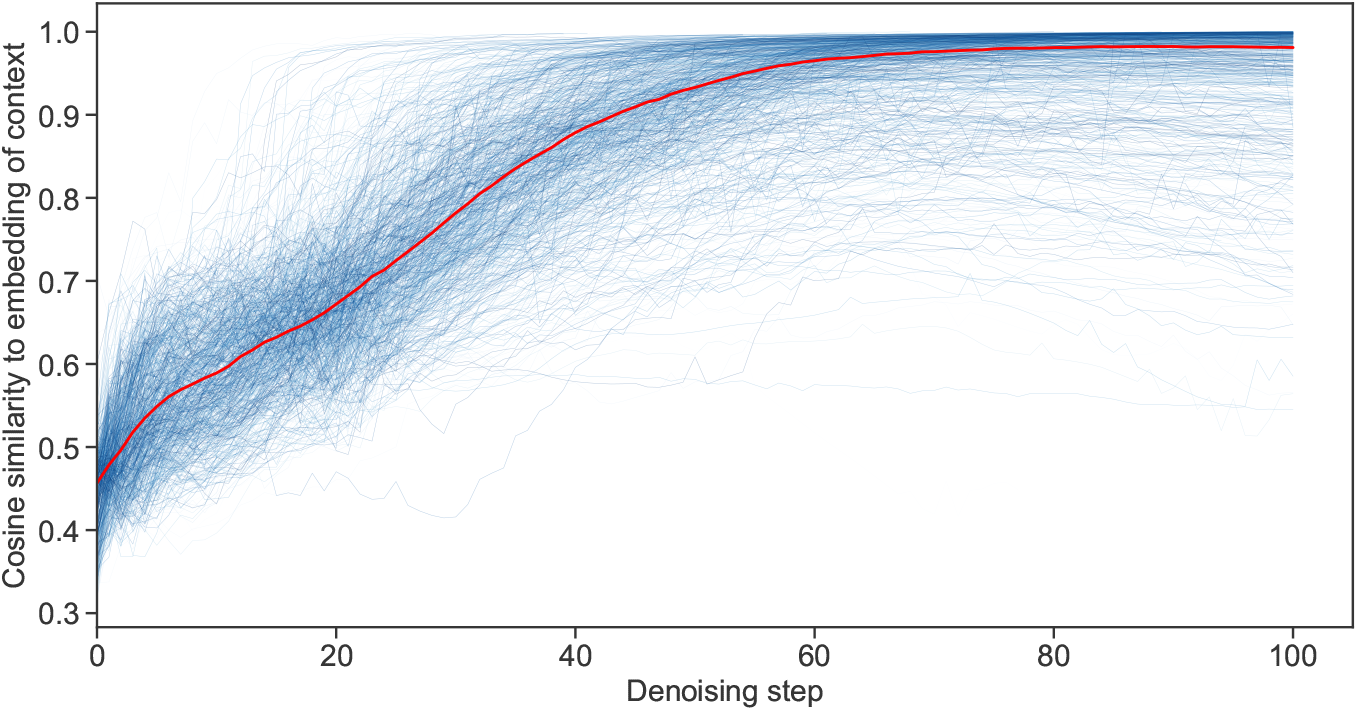
Cosine similarity between the embeddings of generated sequences and those of the context sequences used to condition their generation. Each trajectory is associated to a sequence generated starting using a context sequence from one of the 1000 test set clusters. Cosine similarities were computed at each denoising step and are shown versus denoising step number. The red curve shows the median values. We generated sequences using RAG-ESM (165M), and computed the embeddings as the output of the last layer of the pre-trained ESM2 (650M), so that the similarity measure is not biased by the model used for the generation.

Finally, in Table S6, we compare different strategies for generating novel sequences by denoising a sequence of mask tokens using RAG-ESM, conditioned on context sequences sampled from the test-set clusters. The generation and scoring procedure is the same as in Table 1. However, here, we do not filter sequences based on ESM2 (650M) pseudo-perplexity. Indeed, some generation methods (e.g. without error correction) did not generate enough sequences that passed the perplexity filter.

Table S6 shows that applying Error Correction (EC) during denoising substantially improves the quality of the generated sequences across multiple metrics. Interestingly, delaying EC until halfway through the denoising process (EC@50) yields similar sequence quality compared to applying EC from the start, but results in higher diversity, as measured by the Hamming distance to the context sequence. This suggests that allowing the model to freely sample early residues introduces beneficial stochasticity, and that incorporating EC in the final stage is sufficient to ensure coherence of the generated sequences. We also evaluate a variant that prioritizes the sampling of residues from positions with low entropy logit distributions (i.e. positions the model is most confident about). This method (EC@50 + Ent) slightly decreases generation performance. Therefore, we choose EC@50 as our standard sampling method in Table 1.

**Table S6:** Different generation methods. As in Table 1, we report the median and standard deviation of the distribution of quality scores for generated and natural sequences from 1000 different clusters in the test set. However, contrary to Table 1, sequences are not filtered using ESM2 (650M) pseudo-perplexity. HMMER scores are normalized by dividing each of them by the score of the corresponding context sequence. For natural sequences, Hamming distances to the context are computed for randomly sampled sequences, either from the same cluster as the context (“Homolog”) or from a random different cluster (“Random”). “EC” stands for Error Correction (“EC@50” meaning that the EC process starts from the 50th iteration); “Ent” stands for low-entropy sampling. *↑* (resp. *↓*) indicates that higher (resp. lower) scores are better.

| Score | Generated |  |  |  | Natural |  |
| --- | --- | --- | --- | --- | --- | --- |
|  | No EC | With EC | EC@50 | EC@50 + Ent | Homolog | Random |
| ESMFold pLDDT (↑) | $0.67 \pm 0.15$ | $0.75 \pm 0.14$ | $0.72 \pm 0.15$ | $0.71 \pm 0.17$ | $0.81 \pm 0.16$ | |
| scPerplexity (↓) | $2.83 \pm 0.53$ | $2.62 \pm 0.53$ | $2.67 \pm 0.52$ | $2.76 \pm 0.56$ | $2.58 \pm 0.49$ | |
| ESM2 (650M) perplexity (↓) | $8.50 \pm 2.59$ | $6.32 \pm 2.81$ | $6.69 \pm 2.66$ | $7.17 \pm 2.73$ | $6.06 \pm 3.00$ | |
| RMSD with context (↓) | $5.81 \pm 5.72$ | $4.47 \pm 5.62$ | $5.18 \pm 5.62$ | $4.57 \pm 6.8$ | $9.84 \pm 5.81$ | $11.71 \pm 4.71$ |
| TMscore with context (↑) | $0.42 \pm 0.25$ | $0.48 \pm 0.28$ | $0.44 \pm 0.27$ | $0.47 \pm 0.31$ | $0.19 \pm 0.14$ | $0.14 \pm 0.05$ |
| HMMER (↑) | $0.62 \pm 0.49$ | $0.91 \pm 0.68$ | $0.83 \pm 0.55$ | $0.74 \pm 0.67$ | $0.92 \pm 0.66$ | $-0.01 \pm 0.1$ |
| Hamming to context ($-$) | $0.56 \pm 0.16$ | $0.42 \pm 0.25$ | $0.51 \pm 0.21$ | $0.53 \pm 0.23$ | $0.54 \pm 0.17$ | $0.77 \pm 0.05$ |

### D Motif scaffolding: robustness of model comparisons

**Table S7:** Performance on the motif-scaffolding task. Same as Table 3, except that here the performance of DPLM is the one reported in (Wang et al., 2024) (on just 16 targets).

| PDB ID | Sequence-based |  | Alignment-based | Structure-based |
| --- | --- | --- | --- | --- |
|  | RAG-ESM (165M) | DPLM (650M) | EvoDiff-MSA | RFDiffusion |
| 1bcf | **0.99** | 0.00 | 0.84 | 1.00 |
| 1prw | 0.43 | **0.83** | 0.07 | 0.08 |
| 1qjg | 0.01 | 0.00 | **0.11** | 0.00 |
| 1ycr | 0.05 | **0.38** | 0.00 | 0.74 |
| 2kl8 | 0.01 | **0.08** | 0.03 | 0.88 |
| 3ixt | 0.11 | **0.17** | 0.00 | 0.25 |
| 4jhw | 0.00 | 0.00 | 0.00 | 0.00 |
| 4zyp | 0.00 | 0.00 | 0.00 | 0.40 |
| 5ius | **0.25** | 0.00 | 0.00 | 0.02 |
| 5tpn | 0.00 | 0.00 | 0.00 | 0.61 |
| 5trv | **0.13** | 0.00 | 0.06 | 0.22 |
| 5wn9 | 0.00 | 0.00 | 0.00 | 0.00 |
| 5yui | **0.35** | 0.00 | 0.33 | 0.00 |
| 6e6r | 0.36 | **0.94** | 0.73 | 0.71 |
| 6exz | 0.31 | 0.00 | **0.81** | 0.42 |
| 7mrx | 0.38 | 0.31 | **0.57** | 0.07 |
| Pass rate | **12/16** | 6/16 | 9/16 | 12/16 |
| Mean succ. rate | 0.21 | 0.17 | **0.22** | 0.34 |
| Median succ. rate | **0.12** | 0.00 | 0.05 | 0.24 |

**Table S8:**
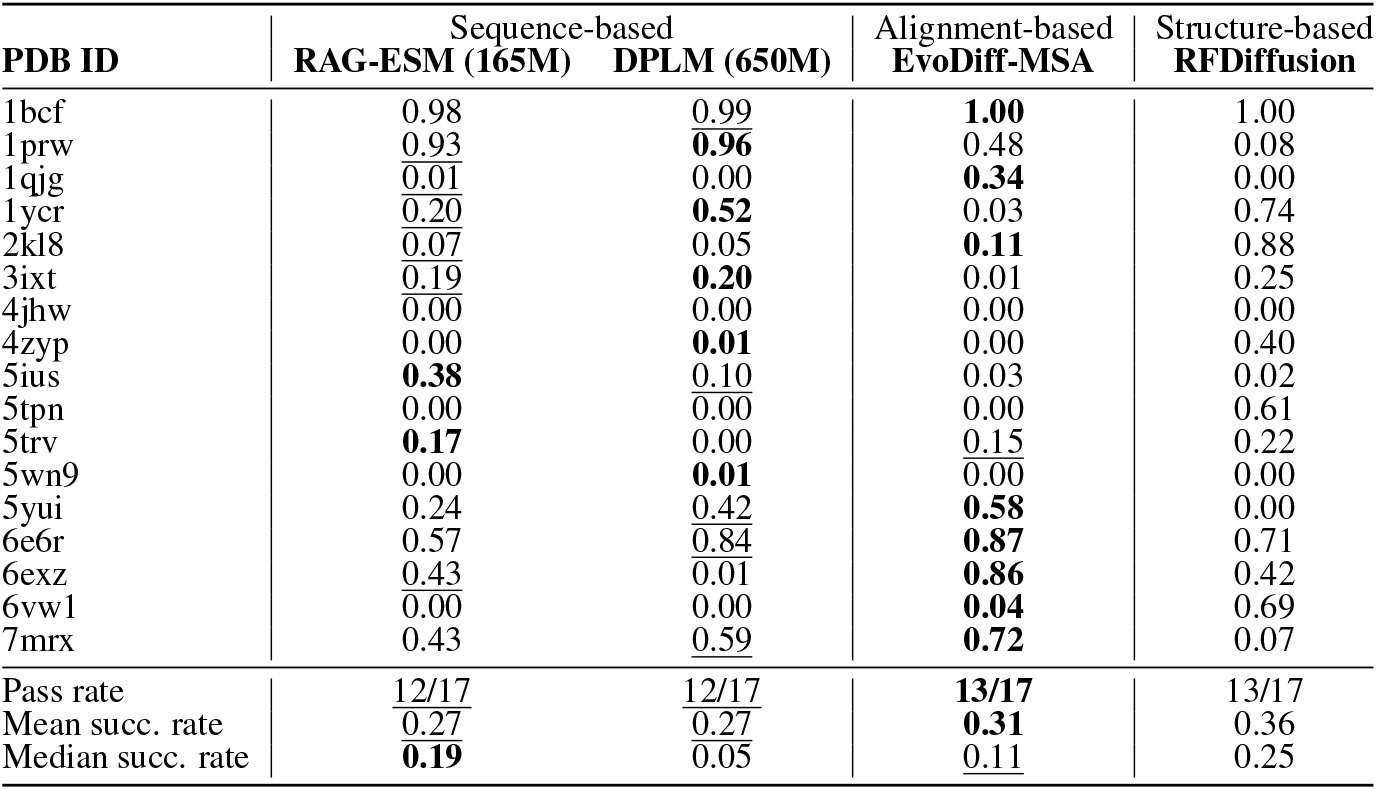
Performance on the motif-scaffolding task. Same as Table 3, except that OmegaFold is used instead of ESMFold as the structure prediction method.

